# The Topological Properties of the Protein Universe

**DOI:** 10.1101/2023.09.25.559443

**Authors:** Christian D. Madsen, Agnese Barbensi, Stephen Y. Zhang, Lucy Ham, Alessia David, Douglas E.V. Pires, Michael P.H. Stumpf

## Abstract

Deep learning methods have revolutionized our ability to predict protein structures, allowing us a glimpse into the entire protein universe. As a result, our understanding of how protein structure drives function is now lagging behind our ability to determine and predict protein structure. Here, we describe how topology, the branch of mathematics concerned with qualitative properties of spatial structures, provides a lens through which we can identify fundamental organizing features across the known protein universe. We identify topological determinants that capture global features of the protein universe, such as domain architecture and binding sites. Additionally, our analysis also identified highly specific properties, so-called topological generators, that can be used to provide deeper insights into protein structure-function and evolutionary relationships. We used our approach to determine structural, functional and disease consequences of mutations, explain differences in properties of proteins in mesophiles and thermophiles, and the likely structural and functional consequences of polymorphisms in a protein. Overall, we present a practical methodology for mapping the topology of the known protein universe at scale.

## Introduction

Proteins are the executors of cellular function, and the main building blocks of cellular structure. Each protein within the protein universe, defined as a collection of all proteins from all organisms, has a three-dimensional (3D) shape (or an ensemble of 3D shapes for a subset of proteins that contain regions of intrinsic disorder), usually referred to as protein structure. One of the main principles of protein science is that shape of the protein determines its function; therefore, determination of the protein structure has been the core interest in molecular biology research over the last seven decades. Collectively, structural biology, structural bioinformatics, and, more recently, deep learning approaches have jointly amassed vast amounts of exquisitely detailed data (*1–4*). AlphaFold2 represents the latest breakthrough in this area. AlphaFold2 is a deep learning model for predicting protein structures that has outperformed accuracy and volume of other protein structure predictions methods. Currently, the AlphaFold2 database contains models for more than 214 million unique proteins across all kingdoms of life, thus covering almost the entire protein universe. Direct analyses of this many structures has thus far been impossible. Capitalizing on the vast computational advances in sequence alignments, (e.g. MMseqs2 (*5*)) it is now possible to use sequence information as a guiding principle for the analysis of the AlphaFold2 database. This allows, for example, sequence-based clustering of structures (*6*), or sequence-based clustering followed by structural alignment of a subset of AlphaFold2 structures (*7*).

However, the majority of commonly used tools for analyzing protein structures and extracting comprehensive and overarching principles governing protein structure and function have been developed to handle much smaller datasets (with notable exceptions, c.f. (*8*)), and to our knowledge, no tool has yet been applied in a structural analysis of the entire AlphaFold2 database. This highlights the need for developing new methods that can be applied for such analysis to unravel the organizing principles of the protein universe.

Protein structures can be described in terms of topology, a powerful framework for understanding connectivity and arrangement of secondary structural elements (e.g. α-helices, β-strands and β-sheets) within a protein (*9*). Mapping of these secondary structural elements and their relationships provides a reductionist view of complex 3D structures of the proteins, and represents a powerful strategy for identifying recurring motifs, spatial arrangements and functional regions within proteins from different organisms and/or protein families (*10, 11*). Therefore, analysis of protein topological features is a cornerstone of protein science that is often used to understand protein structure-function relationships, deduce evolutionary relationships, and engineer proteins with novel functions.

In mathematics, topology is the field that focuses on qualitative features of spatial structures, using algebraic tools. Qualitative features of spatial structures include: connectedness, and holes or voids (*12*). Topology considers any two structures as identical if they can be turned into one another by stretching, twisting, bending, but not cutting or gluing. Thus, the advantage of the topological perspective is that it allows identification of features that are not strongly dependent on the (spatial or temporal) scale at which data are interrogated. In terms of proteins, two proteins that bind the same ligand through similar interactions and similar pockets can be regarded as topologically equivalent, irrespective of their size or detailed global tertiary structure. Therefore, using mathematical topology formalism to analyze protein structures could enable detection of hidden (or latent) structures in complex multidimensional data (*13*).

This might seem vague and unhelpfully general, but the topological perspective has advantages in many settings. Two examples come from physics, and, more recently and pertinently in the present context, topological data analysis (TDA). In physics, Morse theory and Floer homology give exquisite structures to the laws of quantum field theory and cosmology (*14–16*). Recently, TDA emerged as a new approach in mathematical topology (*i*.*e*. topology) (*17, 18*). The essence of TDA lies in analyzing the shape of data using algebraic concepts. The most effective approach to do that is persistent homology (PH) (*19, 20*), a computational tool that transforms scattered points into a sequence of revealing shapes, to identify the system’s features that persist across different scales. When applied to spatial objects (*e*.*g*. protein structures), this corresponds to analyzing how the system’s shape evolves, as its data points become increasingly more spatially extended, overlap and create changing patterns. Thus, PH tracks topological features as they appear and vanish over the course of this spatial filtration, and uses persistence, the measure of how long the feature exists, to distinguish robust signal from noise as the longer a feature persists, the more reliably it captures a feature of the data (*13, 20*). The collection of these features, together with their persistence values, are used as a descriptor of the underlying system and have been proved extremely effective for clustering, parameter inference, or pattern detection in natural and physical systems (*21–24*).

Here, we developed, optimized and implemented a PH-based TDA method to analyze all 214 million structures predicted by AlphaFold2 (*2, 25*). We used this approach to statistically derive organizing principles, topology-function relationships, and obtain a topological “tour guide” to the vast AlphaFold2 resource. In this manner, we address the key need for the field and present a systematic strategy for analyzing the currently largest protein structure dataset in a way that yields insights into structure-function relationships and protein evolution at unprecedented scale.

## Results

### Developing a pipeline for topological analysis of protein universe reveals its topological richness

A recent advance in PH is the ability to efficiently determine “homology generator” (*26*) and to analyze them systematically (*27*). These topology generators pinpoint the specific aspects and regions in the data that are responsible for the creation of topological features. At the level of a single protein, topology generators may reveal groups of highly interacting amino-acids that form higher-order structural features, *e*.*g*. specific conformations (*28*), or entanglement in knotted proteins (*29*). Here, we extended this methodology to analyze more than 214 million protein structures available in the AlphaFold2 database. In order to be able to handle the unprecedentedly large set of topology generators, we developed a feasible process for bulk persistent homology calculation, and to improve memory requirements. The subsequent analysis of the topological output follows the approach developed in (*27, 29*) and by the pipeline in Fig. 1B and Supplemental Fig. 3. As can be seen, in Step 1, we modelled each protein structure via the α-carbon atoms to generate the point cloud representation of the structures. The point cloud representation has the advantage of reducing the complex 3D shape into single points in the (x,y,z) coordinate space for each given residue. The point cloud was used as an input for PH pipeline to compute persistent diagrams and topology generators that provided information about persistence (signal strength or relative relevance/contribution) of each topological feature (in dimensions 1 and 2, *i*.*e*. loops and voids) and interpretation of abstract topological information as local features of the data, respectively (Step 2, Fig. 1B). Thus, the output of this step were topological features, together with their persistence, with each amino acid having the potential to contribute to several, distinct topological features, with different persistence values (Step 3). To understand how important a single region is in affecting the topology of the protein, we computed the point-wise “topological influence score” (TIF), which provided a ranking of amino-acids based on the persistence of their connections (Step 4). TIFs are computed as normalized centrality values on the network of topology generators (*27*), and the TIF values are higher for residues colocated in significant topology generators, see also Supplementary Section 1.Collectively, these steps required circa 66 hours of computational time, performed on Oracle Cloud Compute (see Supplementary Section 3). These computations yielded more than 9.85 terabytes of topological data, and mapped the topology of the currently known protein universe, which we have made freely available online (see Supplementary Section 3).

**Figure 1.**
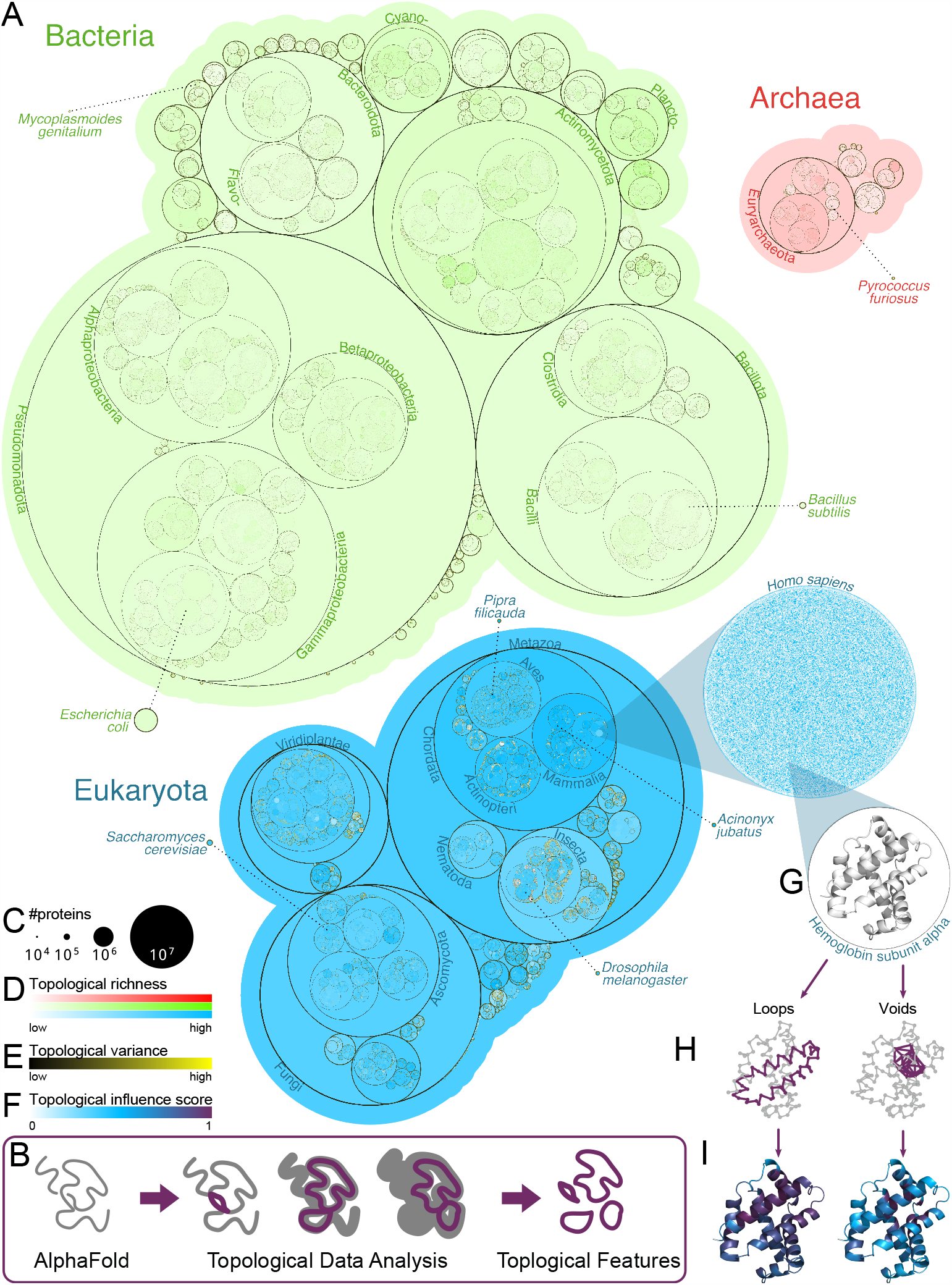
The protein universe is topologically rich. The 214M AlphaFold2 protein structures, organized in species and plotted as a *tree of life* (A). Their topological analysis (sketched in the purple box, B) reveals an intricate variety of features and a remarkable complexity across the evolutionary tree. The tree is constructed using a circle packing plot. The area of each circle is proportional to the number of proteins grouped in it (C). Circles are coloured according to topological richness (D), with boundaries coloured by topological variance (E). Zooming into humans, each protein is shown as a dot, with colour-darkness proportional to its topological richness. Hemoglobin (G) is plotted showing its most persistent one-dimensional (left, a loop) and two-dimensional (right, a void) topological features (H), and below with amino-acids colored by their topological influence score (I), with a white-blue-purple scale (F).

To examine the resulting data in the broadest and most general context, we used it to construct the topological tree of life (Fig. 1A). The tree was constructed using a circle packing plot, with the area of the circle corresponding to the number of AlphaFold2 predicted structures available for each species. Next, we connected and ranked each genus and related them one genus at the time, so that the area of higher ranks is only approximately representative of the number of structures (Fig. 1C). The three domains of life, bacteria, archaea and eukaryotes, are all well represented in the topological tree of life, and include organisms with vastly varying sizes of their proteomes. Furthermore, we were able to map the topological richness for each protein, each organism and across domains (areas of low richness in light colors, areas of high richness in dark colors, (Fig. 1D). Topological richness is the measure of how many unique, persistent topological features each protein has, averaged across all proteins and normalized by number of residues. We observed that comparatively speaking, bacterial and archaeal proteins exhibited lower topological richness, whereas eukaryotes exhibited several areas of heightened richness, especially within the mammalian class.

A couple of notable highlights among species include *Acinonyx jubatus* (cheetahs) and *Pipra filicauda* (wire-tailed manakin), while humans are outliers among other mammals, in terms of their low value. It might seem surprising that humans show a lower richness than other species in their class. However, similarly to the case of gene count, which were found to be unexpectedly low for humans, this indicates that topology is just one among many ways to measure complexity. In this specific case, human complexity at the protein level is more likely to arise from intricate layered regulation (*30*) and the complexity of the protein-protein interaction network (*31*).

Furthermore, we mapped the topological variance (Fig. 1E), which can be taken as a measure of evolutionary robustness of topological characteristics. The topological variance is computed as the variance of the number of 1-dimensional topological features in a given circle, normalized by the number of proteins in the circle. The variance is shown in the figure as the outline of disks, using a black-yellow color-code. Similarly to richness, topological variance is higher for eukaryotes than for Bacteria and Archaea. This is particularly evident in insects, especially at the species level, and suggests an increased diversity in their topological features. On the other hand, when variance is consistently low across ranks (as for Bacteria), this can be interpreted as topological complexity being preserved through evolution.

Lastly, we mapped TIFs on each individual protein, which gives information at the residue level (Fig. 1F and I). In each protein structure, TIF values quantify how topologically important each residue is, which, in turn, leads us to identify structurally significant regions, and locations for potentially damaging mutations, as we show in Section Topological analysis detects protein regions enriched for disease-associated mutations.

Beyond the topological tree of life view, we can zoom in on individual proteins such as human hemoglobin subunit alpha (Fig. 1G), where our analysis identified its most persistent loop and void (Fig. 1H), and how they influence the topology as measured via TIFs (Fig. 1I). Taken together, our new method offers an unprecedented tool for analyzing topology of the protein universe. Applying our pipeline to the whole AlphaFold2, a database with close to a quarter of a billion protein structures, reveals both the intricacies and variety of topological features across the tree of life.

### Topological analysis of the protein universe enables nuanced protein structure analysis

We have recently shown that PH can be analyzed using network theory, which reveals further relationships between topological features (*27*). To extract further insights from our topological map of the protein universe, we interpreted protein topology as a network, where edges are defined by loops (dimension 1) and voids (dimension 2). In this framework, the intensity and overlaps of these connections induce a grouping of amino acids into units, which we call “topological clusters” (Fig.). This approach allowed us to capture global structural properties of the protein universe that detected characteristics beyond traditional strategies for protein structure analysis.

For example, we observed that topological clusters of dimension 1 (loops) are closely associated with protein domains as defined by e.g. the CATH Protein Structure Classification Database (*32*). We illustrated this by examining more closely the relationship between CATH domains (Fig. A) and topological clusters (Fig. B) of a protein kinase (UniProt (*33*) ID Q4DF08) as a representative example. We noted that in this case, as well as many others, topological clusters of dimension 1 capture the essence of CATH domain classification; more specifically, we noted that here a single CATH domain is partitioned into multiple topological clusters. In other words, the topological analysis refines on the resolution provided by domain assignments. We used homogeneity score to quantify whether the topological clusters of dimension 1 provide an exact subdivision of CATH domains (score = 1), or whether the two partitions are completely unrelated (score = 0). We analyzed the 38,171 AlphaFold2 structural predictions, representing different protein families, domains, and organisms (Supplementary Table 2), which correspond to all non-redundant, high-confidence, AlphaFold2 predictions, containing at least two distinct identified CATH domains (*25, 32*) (see also Supplementary Section 4). As seen in Fig. C (see also Supplemental Fig. 8, showing the same computation, but including redundant structures), the vast majority of topological clusters belong to a single domain; thus, the topological analysis refines on the resolution provided by domain assignments, revealing that many domains are formed by distinct topological features. This may have important implications for evolutionary analysis as well as protein engineering efforts, given that work in these areas very often use protein domains as basic units of analysis. Our results suggest that, for the majority of proteins, mathematical topology is consistent with and refines into more nuanced features known protein domains cataloged in CATH and similar databases.

We can further speculate that topological clusters could identify specific folding units. Folding intermediates are difficult to obtain, but experimental evidence for partial folds apomyoglobin forming within micro-and milliseconds exists (*34–36*). In this case, the 1-dimensional topological analysis captures the initial folding core (the blue cluster in Fig. F). While this is encouraging preliminary evidence that the topological perspective can augment the analysis of protein folding, as noted elsewhere (*40*), AlphaFold does not provide the structural ensembles necessary to shed light on protein folding dynamics.

Unlike dimension 1 topological features (loops) that could inform on substructures, dimension 2 (voids) may be associated with binding sites. To examine if this was the case, we examined the distances between the clusters of voids and the binding sites as defined in the Mechanism and Catalytic site Atlas (M-CSA) dataset (*37*). Our analysis included 866 AlphaFold2 predicted protein structures (862 predicted with high confidence), representing a broad range of enzyme families and other proteins known to engage ligands (Supplementary Table 2). These structures were obtained by mapping to UniProt all 1033 RCSB Protein Data Bank (RCSB PDB) (*38*) entries of experimental structures having M-CSA annotated sites, and then by selecting those corresponding to high-confidence AlphaFold2 predictions. We mapped the distance in terms of number of residues between the void boundary and the binding site and observed that some 70% of binding sites are either immediately at the boundary of a void or one amino acid away (Fig. D, see also Supplemental Fig. 7, showing the same computation, but including low-confidence predictions). Again, this makes sense from a structural perspective, as binding sites must correspond to areas of accessibility and flexibility, and our topological analysis allows the detection of such sites across 214 million predicted structures. Thus, mapping voids has the potential to identify cryptic and/or unknown binding sites within the protein universe.

Despite its remarkable accuracy, AlphaFold2 predictions can sometime fail to fully capture the structural complexity of certain proteins (*39–41*). One of PH’s most powerful features is its robustness: input data differing by small perturbations will have similar topological fingerprints (*42*). Thus, it is reasonable to assume that our topological analysis will be blind to possible misinterpretation of local 3D conformations. To assess this, we compared the output of our pipeline for experimental structures cataloged in RCSB PDB with their AlphaFold2 counterpart, results are shown in Fig. E. We quantified the discrepancy between topological features in the experimental and predicted datasets by looking at TIFs in dimension 1 and 2; as shown by the bar-plot, per residue values are highly correlated in both cases, ensuring that the topological analysis is transferable from simulations to experiments.

Taken together, we demonstrate the value of topology in identifying features of protein structural organization.

### Topological comparison of thermophilic and mesophilic proteins

How thermophilic proteins achieve stability while maintaining functionality remains a heavily debated question in protein science and structural biology. Different factors, such as differences in hydrophobicity, secondary structure, ion-pairing, hydrogen bonds, and numbers and sizes of cavities, have been proposed as key determinants of thermophilic protein stability and function (*43*). However, lack of statistical power and the need to correct meticulously for potentially confounding factors has impeded progress. Given the wealth of structural information generated by AlphaFold2 and the robust nature of topology, we hypothesized that we can detect topological differences between – even structurally very similar – thermophilic and mesophilic proteins, and that these differences may provide insights into how thermophilic proteins maintain their structure and function. Finding such differences is especially challenging as, across different organisms, specific enzymes often present highly similar, almost super-imposable structures, as it happens for example in the case of Glucose-6-phosphate 1-dehydrogenase, shown in Fig. 3A in *E. coli* (mesophile) and *M. thermoacetica* (thermophile).

**Figure 2.**
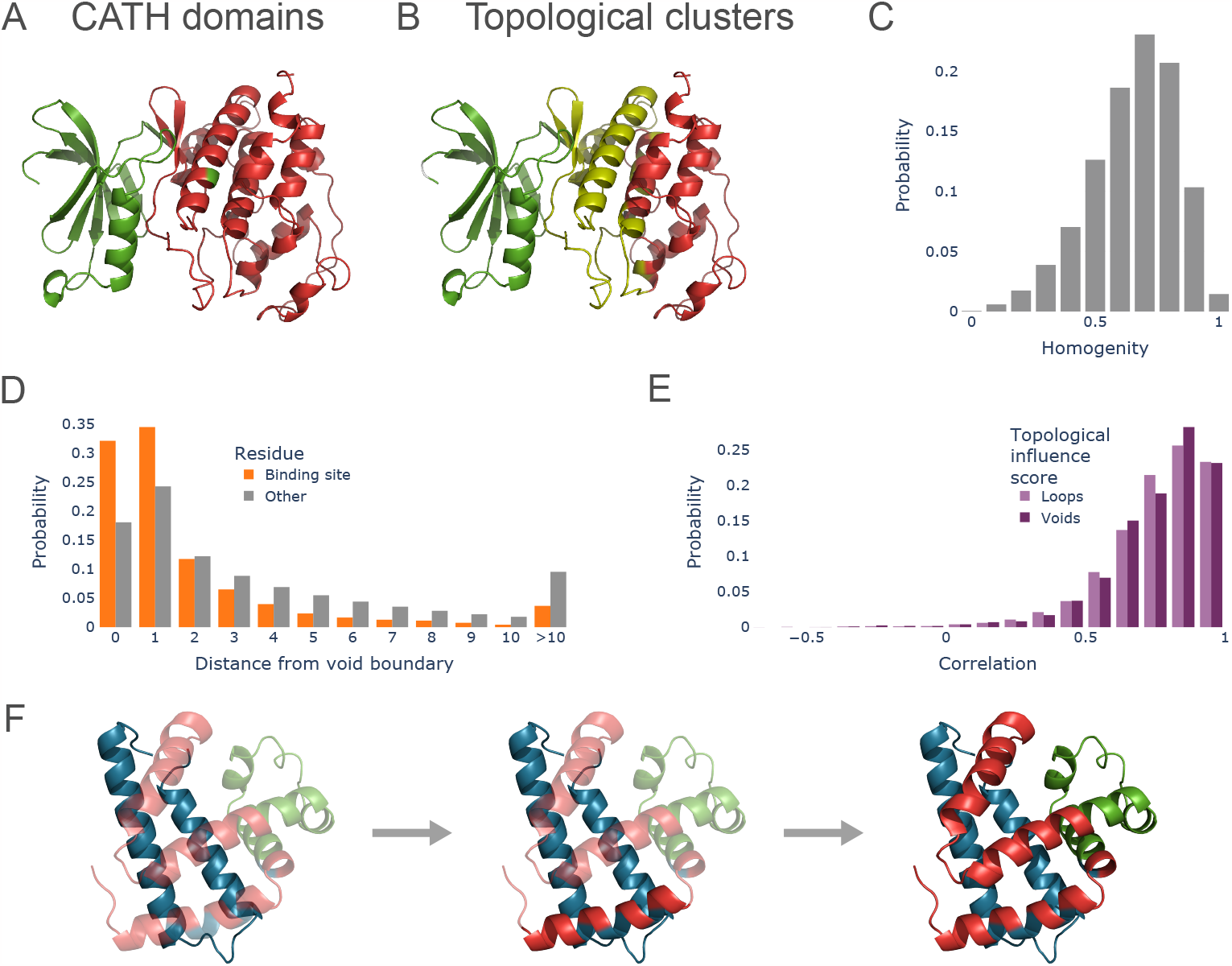
Topology provides organizing principles for predicted protein structures. Topological clusters in dimension 1 often provide a refinement of protein domains. As an example, we see protein kinase, colored by its two CATH domains (A) and topological clusters (B). The homogeneity score (C) can be used to check if a topological cluster contains only residues belonging to a single domain, with 1 corresponding to a perfect sub-partition. The bar plot shows the distribution of homogeneity scores for a set of 38,171 non-redundant AlphaFold2 predictions with identified CATH domains. (D) The distribution of distances (in number of residues) of residues between binding sites from boundaries of 2-dimensional topological clusters. These boundary points are enriched for binding sites from the Mechanism and Catalytic Site Atlas (M-CSA) dataset. (E) The topological analysis is robust to small perturbations. The bar plot shows the distribution of correlation coefficients between the topological influence scores between AlphaFold2 predicted structures and their experimentally solved counterparts. The high values show that topological features tend to correspond and to interest the same residues. (F) Topological clusters for horse apomyoglobin (UniProt accession P68082) differentiated by color. Folding events based on experimental evidence is indicated by transparency, where fully opaque sections are formed.

**Figure 3.**
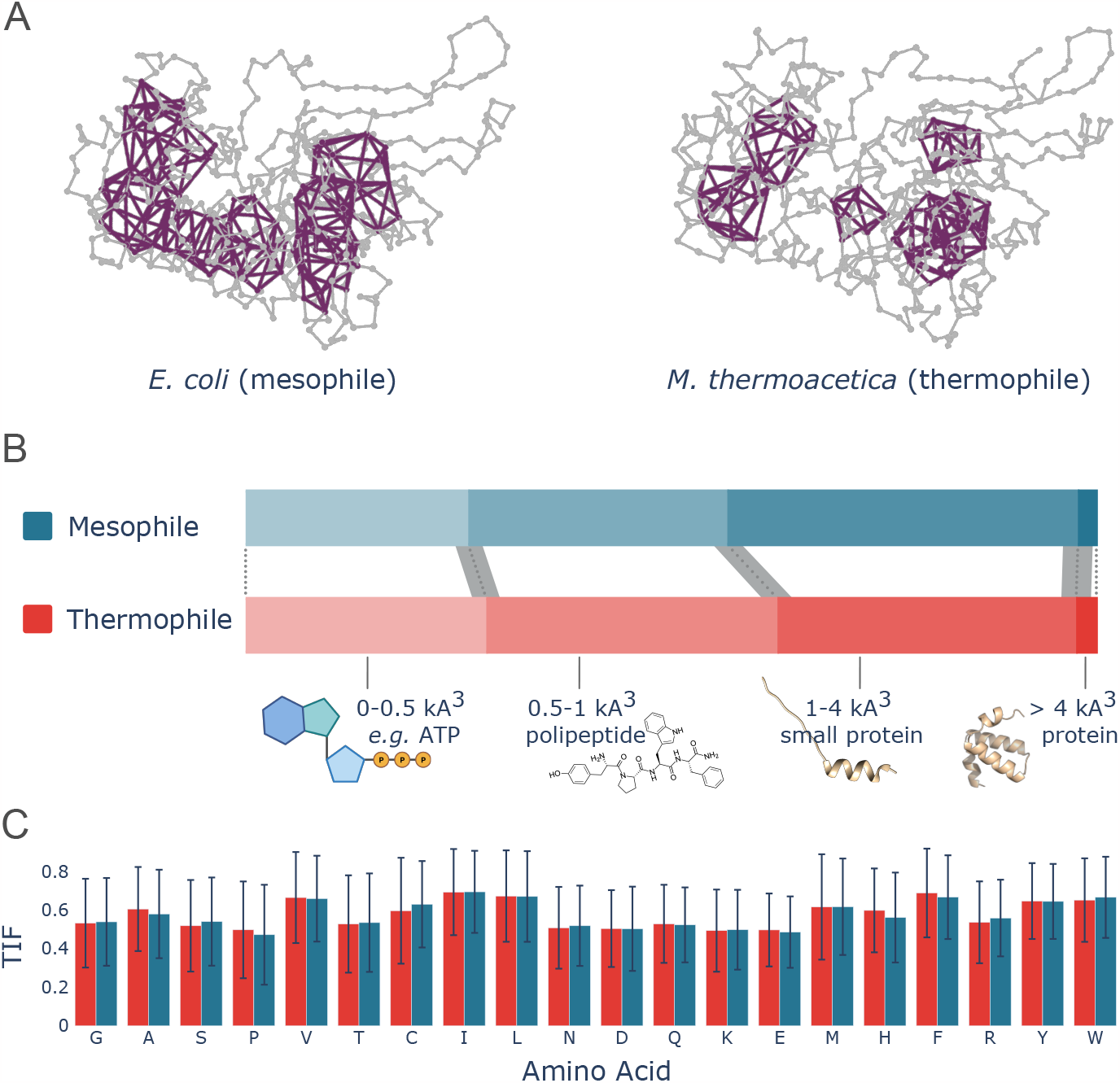
Thermophilic and mesophilic proteins are topologically different. Highly per-sistent topological features in dimension 2 identify voids in protein structures. Voids in ther-mophilic organisms are in general smaller in volume than in their mesophilic counterparts, as shown here by Glucose-6-phosphate 1-dehydrogenase (A). The observed pattern is consistent across enzymes from 10 different EC numbers (B) and robust. Error bands are computed as the standard deviation of the different ratios after sampling 1,000 voids among thermophiles and mesophiles. Further, the result is not affected by amino acid frequencies, or TIF distributions, as these are identical (C).

To address this, we selected 10 different Enzyme Commission (EC) numbers based on their relevance in biotechnology (see Supplemental Table 2 and Supplemental Fig. 9, 10 and 12 for details). The selected enzymes covered 30 thermophilic and 8 mesophilic organisms, for a total of 1,656 high confidence AlphaFold2 predictions. We compared topological features of dimension 2 – *i*.*e*. voids – in mesophiles (blue) and thermophiles (red) (Fig. 3B). For this analysis, we chose to focus on voids because we were interested in understanding whether more compact topological features could be associated with high-temperature preferences.

Additionally, we focused on comparing orthologous proteins with matching amino acid sequence length to minimize potential compounding effects of variable protein sequence length, substrate/binding partner properties and function. We observed that voids in predicted protein structures from thermophilic organisms are smaller and more compact than their mesophilic equivalents (Fig. 3B), and that this difference is statistically significant (Supplemental Fig. 11). Furthermore, the same pattern of decreased void volume is also apparent when considering enzymes by their category (Supplemental Fig. 12). However, this difference in the size of the voids can’t be explained by a simple difference in amino acids composition, as we detect no difference in the amino acid composition of voids for thermophiles and mesophiles (Fig. 3C). Based on these results, we speculate that the topological differences between thermophiles and mesophiles reflect the different thermodynamic pressures experienced by the different organisms in their respective habitats, where binding pockets with larger voids may not provide the correct specificity of binding at higher temperatures, and the adequate thermodynamic stability.

### Topological analysis detects protein regions enriched for disease-associated mutations

Given the fact that protein function depends on protein structure and sequence, we examined whether topological analysis can detect protein regions that are enriched in damaging, disease-associated mutations. Thus, we used a dataset of disease-causing and neutral variants that contains experimental structures of a few hundred wild-type and mutated proteins (*44, 45*). This dataset was previously analyzed to establish the link between damaging mutations and their effect on structures (*44*). As in the above instances, we restricted our analysis to structures predicted with high confidence. For each of the proteins analyzed, we wanted to identify residues that are structurally important, and thus, more likely to accommodate mutations leading to structural damage, and in turn, to the occurrence of disease-associate polymorphisms. TIF values provide a measure of the topological significance of each residue; a natural question is whether a high 1-or 2-dimensional TIF directly estimates the influence on structural stability. Overall, we found that mutations that give rise to structural variants, those that give rise to disease, and those that give rise to disease and structural effects, are more likely to be co-located with topology generators than non-disease causing variants, or polymorphic sites that have no known structural role. Figure 4A and B shows the 3D structures of human ACE2 (top) and HBB (bottom), colored by their per-residue two-dimensional TIFs. On the right-hand side, we see the distribution of 2-dimensional TIFs on residues whose substitution induce polymorphisms that are predicted to be structurally damaged and associated with disease, or neutral (*45*). In these examples, the pattern discussed above is clearly visible. A similar result is observed in other individual proteins (see *e*.*g*. Human Adenylosuccinate lyase, Fig. 4C, and CTFR, Supplemental Fig. 15), in the whole dataset considered (Fig. 4B, and Supplemental Fig. 14), and for 1-dimensional features alike (Supplemental Fig. 13).

**Figure 4.**
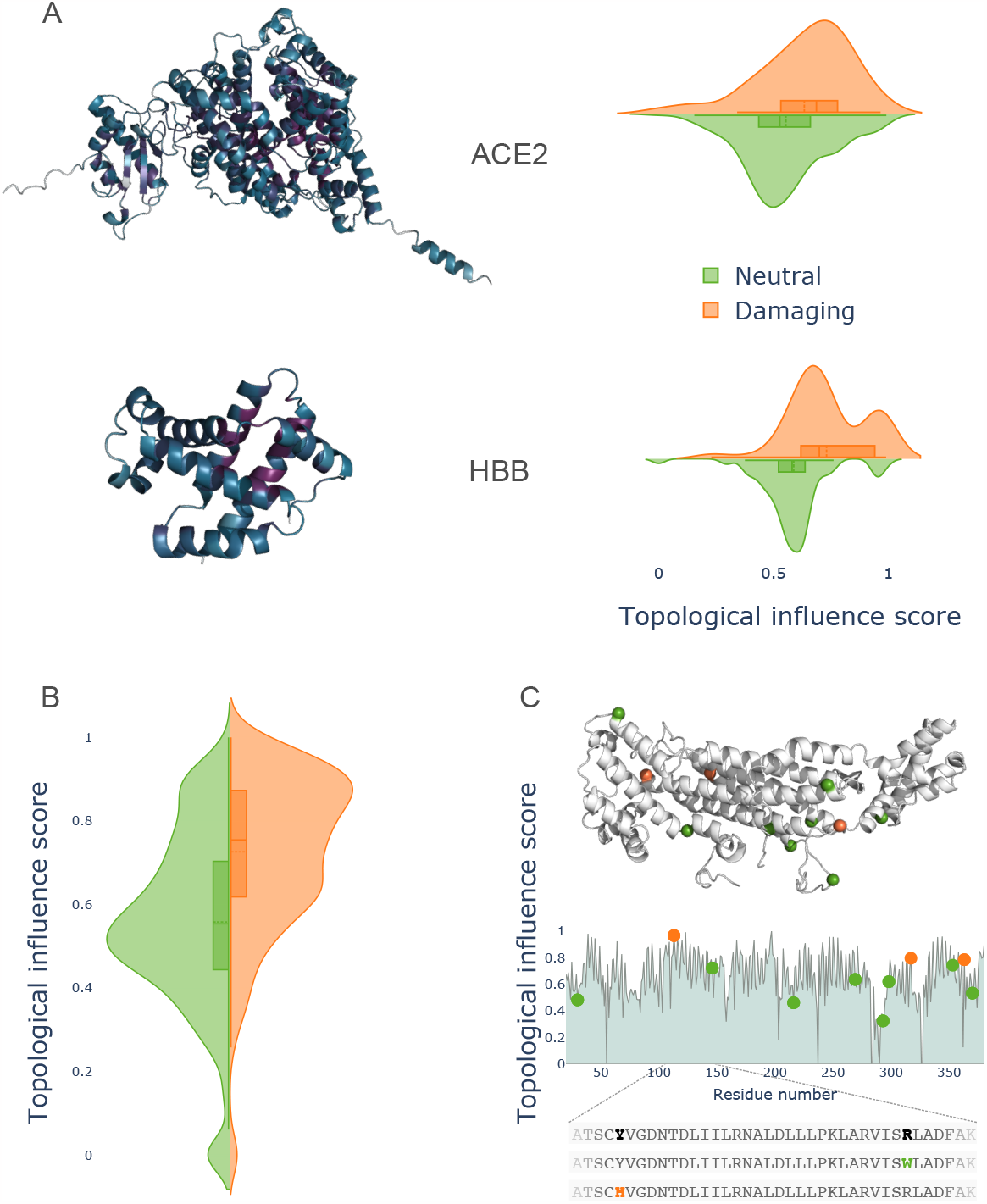
Topological features are enriched in damaging variants. (A) The 3D structures of human ACE2 (top) and HBB (bottom), colored by their per-residue two-dimensional TIFs. Structural analysis of missense variants for these genes predict a number of them to be damaging (*44, 45*). Our topological analysis shows that amino acid substitutions causing structural damage are more likely to happen where the TIF is high, as shown by the violin plots on the right-hand site. This pattern is maintained across a dataset of disease-associated missense variants (B). As a further example, (C) shows human Adenylosuccinate lyase, with its missense variants highlighted on the 3D structure, and a plot of its 2-dimensional TIFs on the bottom.

## Conclusions

In this work we demonstrate that topology can serve as an interpretative tool for the data that is contained in the AlphaFold2 database. To achieve this, we developed a pipeline that can achieve topological analysis of all 214 million predicted protein structures in a time-and cost-effective manner. We showed how the topological information can be used to extract global insights into the features and properties of the protein universe. Furthermore, we described several use case scenarios, including using topology to analyze coarse structural features, such as domains and binding sites, to map distinctions between thermophiles and mesophiles, and examine effects of disease-causing mutations. To enable further work by the broader community, we provide access to the one and two-dimensional persistence diagrams, topological features, and TIFs (per residue) as the online resource with the total resource amounts to approximately 20 TB.

Overall, we demonstrated that topology allows us to make sense of the vast amounts of protein structural data at the required scale. Importantly, our analysis was done using solely structural data on positions of *Cα* provided by AlphaFold2 (and the PDB for validation) without any biological knowledge or sequence information. Thus, going forward, incorporating additional pieces of information, such as the biophysical and biochemical properties of the amino acids and their three-dimensional arrangement, may capture a host of additional factors that influence protein function. Nonetheless, our work highlights that topology adds an additional set of features for function prediction, or an additional dimension to the biophysical analysis of protein structure. Although topology may not be enough to rationalize protein function, we are confident that topology offers a natural and direct route for making sense of the wealth of data in AlphaFold2; and that the topological information generated here will aid the functional and evolutionary analysis of the molecular machinery of life.

## Acknowledgments

CDM gratefully acknowledges funding through a University of Melbourne PhD studentship. AB gratefully acknowledges funding through a MACSYS Centre Development initiative from the School of Mathematics & Statistics, the Faculty of Science and the Deputy Vice Chancellor Research, University of Melbourne. SYZ gratefully acknowledge funding through a University of Melbourne PhD studentship. MPHS is funded through the University of Melbourne DRM initiative, through an ARC Laureate Fellowship (FL220100005) and acknowledges financial support from the Volkswagen Foundation through a “Life?” program grant. LH is funded through the University of Melbourne DRM initiative. AD gratefully acknoledges funding through Wellcome Trust grant 218242/Z/19/Z. DEVP received funding from an *Oracle for Research* Grant. The authors wish to thank Ellen Leffler for helpful discussions.

