## Supplementary Information for "The Topological Properties of the Protein Universe"

### 1. Pipeline and computations

1.1. **Persistent homology.** Persistent homology [19, 12] is a method in computational topology for analysing the shape of data via topological features. Persistent homology is built on the concepts of *simplicial complexes* and s*simplicial homology* [11]. Intuitively, a simplicial complex is a space constructed by gluing together *simplices* (*i.e.* points, line segments, triangles, and their higher dimensional counterparts), for a formal definition see *e.g.* [11, Ch.2]. Let $PC = \{p_1, \dots, p_n\} \subset \mathbb{R}^n$ be a point cloud, *i.e.* a set of scattered points in the Euclidean space $\mathbb{R}^n$; the shape of $PC$ can be described by constructing a simplicial complex $PC_\varepsilon$ that approximates the connectivity of the points $p_i$ at a given spatial scale $\varepsilon$. Common choices of such a simplicial complex are:

- The Vietoris-Rips complex [12, Ch.III.2] $PC_\varepsilon = VR_\varepsilon(PC)$; this is constructed by adding a $k$-simplex $[v_{i_0}, v_{i_1}, \cdots, v_{i_k}]$ if the distance between all pairs of points in $\{v_{i_0}, v_{i_1}, \cdots, v_{i_k}\}$ is less than $\varepsilon$.
- The Ĉech complex [12, Ch.III.2] $PC_\varepsilon = C_\varepsilon(PC)$; this is constructed as the *nerve complex* [11, Ch.3] of the union of balls of radius $\varepsilon$ centred in $PC$.
- The Alpha complex [12, Ch.III.4] $PC_\varepsilon = \mathrm{A}_\varepsilon(PC)$; this is similar to the Ĉech complex, but has a canonical geometric realisation, and it is a sub-complex of both the Delanauy complex and the Ĉech complex.

Note that for each of these choices, $PC_{\varepsilon_1} \subset PC_{\varepsilon_2}$ whenever $\varepsilon_1 < \varepsilon_2$. More information on these complexes, their differences, and their properties can be found *e.g.* in [12]; see also Figure 1 for one example.

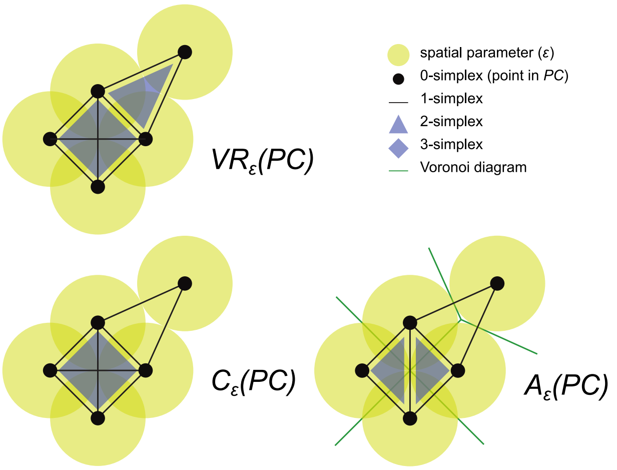


**Supplementary Figure 1.** An example of Vietoris-Rips complex, Ĉech complex and Alpha complex constructed on the same point cloud, for the same parameter $\varepsilon$.

The qualitative features of $PC_\varepsilon$ can be analysed by computing its $k$-dimensional simplicial homology $\mathrm{H}_k(PC_\varepsilon; \mathbb{F}_2)$, where $\mathbb{F}_2$ is the field with two coefficients. For each choice of dimension $k$, $\mathrm{H}_k(PC_\varepsilon;\mathbb{F}_2)$ is a vector space, and its rank corresponds to the number of $k$-dimensional topological feature (called homology classes) of $PC_\varepsilon$. The 0-dimensional homology $\mathrm{H}_0(PC_\varepsilon;\mathbb{F}_2)$ counts the "connected components" (*i.e.* separate pieces) that form $PC_\varepsilon$, while 1 and 2-dimensional homologies $\mathrm{H}_1(PC_\varepsilon;\mathbb{F}_2)$ and $\mathrm{H}_2(PC_\varepsilon;\mathbb{F}_2)$ count loops and voids respectively. For a formal definition of simplicial homology, see *e.g.* [11].

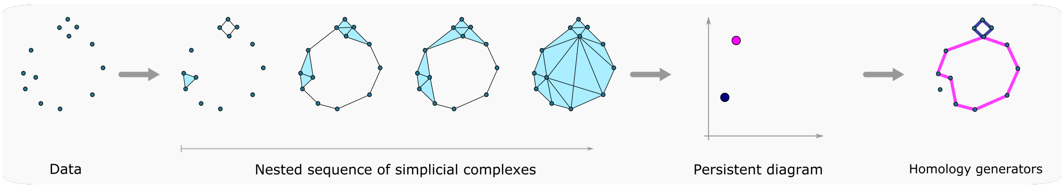


**Supplementary Figure 2.** In the standard persistent homology pipeline, the input is a point cloud $PC = \{p_1, \ldots, p_n\} \subset \mathbb{R}^n$. The first step in the algorithm is to construct a nested sequence of simplicial complexes $PC_{\varepsilon_0} \hookrightarrow PC_{\varepsilon_1} \hookrightarrow \cdots \hookrightarrow PC_{\varepsilon_N}$ that approximate the data at different spatial scales. This depends on a choice of *filtration*; common choices are the Vietoris-Rips, Ĉech and Alpha complexes. The figure shows a schematic of the Vietoris-Rips filtration built on the initial point cloud. The simplicial complexes are then analysed by computing their simplicial homology and corresponding persistent diagrams. The figure shows the output for 1-dimensional features; a total of two loops appear in the filtration. The fact that one is more persistent than the other reveals the *circular* structure of the data. For each homology class in a persistent diagram, it is possible to compute representative cycles. The figure shows a possible choice of generator for each 1-dimensional feature in the persistent diagram.

Persistent homology studies the shape of the initial data $PC$ at different spatial resolutions, by looking at the simplicial complexes $PC_\varepsilon$ for increasing values of $\varepsilon > 0$, see Figure 2. This results in a nested sequence of simplicial complexes

$$PC_{\varepsilon_0} \hookrightarrow PC_{\varepsilon_1} \hookrightarrow \cdots \hookrightarrow PC_{\varepsilon_N}$$

which in turn yields a sequence of vector spaces and maps between them

$$\mathrm{H}_k(PC_{\varepsilon_0};\mathbb{F}_2) \to \mathrm{H}_k(PC_{\varepsilon_1};\mathbb{F}_2) \cdots \to \mathrm{H}_k(PC_{\varepsilon_N};\mathbb{F}_2)$$

called the $k$-dimensional *filtered* homology of $PC$.

We are interested in looking at how topological features evolve in this sequence of simplicial complexes and homology spaces. Thanks to the Structure Theorem [18, Thm 2.1], we can summarise the information contained in each sequence $\mathrm{H}_k(PC_{\varepsilon_0};\mathbb{F}_2) \to \mathrm{H}_k(PC_{\varepsilon_1};\mathbb{F}_2) \cdots \to \mathrm{H}_k(PC_{\varepsilon_N};\mathbb{F}_2)$ as a "persistent diagram" $PD$. This is a finite collection of points $PD = \{(b_i, d_i)\}$, where $b_i$ and $d_i$ are the birth and death scales of the $i^{\text{th}}$ $k$-dimensional feature. The "persistence" of each feature is given by the difference $d - b$, which gives a measure of its significance.

For each homology class, it is possible to compute a "representative" or "generator", that is, a specific set of simplices creating the corresponding homology feature [11]. Homology generators provide an interpretation of the abstract topological information as local, structural features of the data [48, 27, 28, 26].

1.2. **Topological analysis of protein structures.** The topological analysis of the protein universe follows the methodology developed in [28, 26], see Figure 3 for a schematic representation.

**Step 1.** We model each protein structure as the point cloud given by its α-carbon atoms, *i.e.* by the set $PC = \{p_1, \ldots, p_n\}$, where each $p_i = (x_i, y_i, z_i)$ is the triple of the predicted $xyz$-coordinates of its $i^{\text{th}}$ residue.

**Step 2.** We then feed the point cloud $PC = \{p_1, \ldots, p_n\}$ to the persistent homology pipeline, and compute its filtered homology in dimension 1 and 2:

$$\mathrm{H}_1(PC_{\varepsilon_0}; \mathbb{F}_2) \to \mathrm{H}_1(PC_{\varepsilon_1}; \mathbb{F}_2) \cdots \to \mathrm{H}_1(PC_{\varepsilon_N}; \mathbb{F}_2)$$

$$\mathrm{H}_2(PC_{\varepsilon_0}; \mathbb{F}_2) \to \mathrm{H}_2(PC_{\varepsilon_1}; \mathbb{F}_2) \cdots \to \mathrm{H}_2(PC_{\varepsilon_N}; \mathbb{F}_2).$$

From these, we compute the persistent diagrams in dimension 1 and 2.

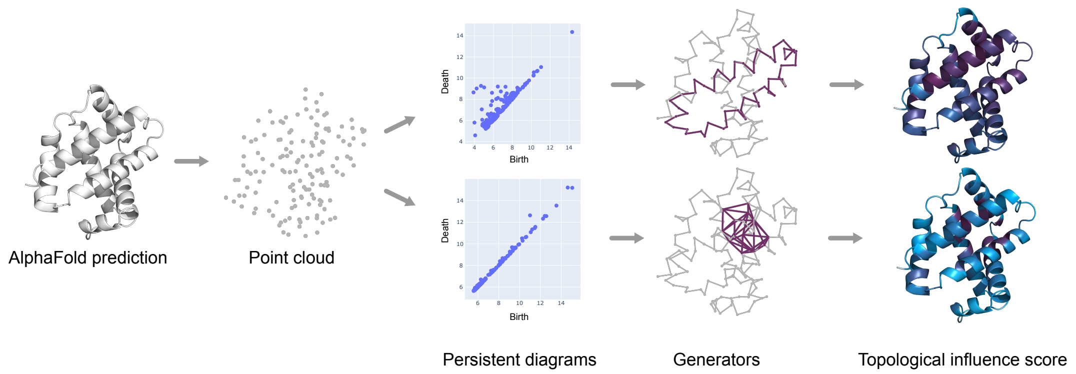


**Supplementary Figure 3.** From a predicted protein structure, we extract the point cloud $PC = \{p_1, \ldots, p_n\}$ given by $xyz$-coordinates of its α-carbon atoms. We then compute the corresponding persistent diagrams in dimension 1 and 2, and a generator for each topological feature. For each dimension considered, the information consisting of persistent diagram, generators, and their persistence is used to compute the point-wise topological influence score (TIF), which ranks residues based on how often they contribute to topological features, and how important these are.

**Step 3.** We compute a representative cycle for each homology class. Note that these correspond to loops and voids appearing in the sequence of simplicial complexes $PC_{\varepsilon_0} \hookrightarrow PC_{\varepsilon_1} \hookrightarrow \cdots \hookrightarrow PC_{\varepsilon_N}$.

**Step 4.** We compute the 1 and 2-dimensional point-wise topological influence score (TIF) of residues in $PC$. This is achieved by first computing centrality values centrality(res) for each residue, as in [26] and using spectral methods developed in [50]. Then, centrality scores are normalised over all the residues in the protein to obtain values in $[0, 1]$:

$$\mathrm{TIF(res)} = \frac{\mathrm{centrality(res)}}{\mathrm{max_r(centrality(r))}}.$$

TIFs provide a ranking of residues based on how often they contribute to topological features (*i.e.* how often they appear in generators) and how persistent these features are.

1.3. **Software.** Persistent diagrams and generators are computed using the Julia software `Ripserer.jl` [25]. Specifically, we use the Alpha filtration to construct the nested simplicial complexes, and the involutive algorithm [25, 47] to compute homology and representatives.

TIFs are computed using the hyperTDA method developed in [26]. Specifically, for each protein structure and dimension considered, we construct the hypergraph having as vertices the residues, and having a (weighted) hyperedge for each generator. Then, we compute node centrality using the software from [50], using the *max* centrality flavour. More details are contained in the hyperTDA paper [26] and corresponding GitHub repository.

Similarly, topological clusters are computed as graph-communities, as explained in [26] and using Python's Louvain module [45].

1.4. **How we handled computations.** Large-scale computations were performed on Oracle Cloud Compute. All computations were performed on a single instance with 160 CPU cores and 1 TB memory. The *compute shape* is named `BM.Standard.A1.160` which is Arm-based Ampere A1 compute (Ampere Altra processor). A 32 TB block storage volume was attached for storage of AlphaFold2's predicted structures as well as general storage, and a separate 32 TB volume for the outputs of our topological analyses. The former was mounted at the project root, and the latter at `data/alphafold/PH/`.

AlphaFold2 structures were downloaded as sharded proteomes according to their bulk download instructions.

Benchmarking was performed on a single large structure (accession "A0A009DWL0") to assess the computational viability and reduce time, cost, and environmental impact (Fig 4A). Only homology dimension one was computed. Julia methods were run multiple times before the recorded run, to remove the impact of compilation. Note that some bars appear to have zero height, since methods in compiled languages such as C++ have significantly lower memory consumption than Julia and Python methods. Considerations to computational cost were also important in terms of the memory usage, as the cluster becomes unstable when the 1 TB is exceeded (Fig 4C). Tools that were benchmarked:

- **`Eirene.jl`:** the initial method used in previous works due to its ability to compute representative cycles.
- **`Eirene.jl mod`:** a modified version of `Eirene.jl`, which was made in an attempt to tailor it to this specific project, however this barely improved time at the cost of increased memory consumption.
- **`giotto-ph`:** a method written in C++ and Python which takes advantage of CPU parallelisation. It was not considered further, as it does not compute representatives.
- **`Gudhi`:** a toolkit with numerous Python modules, however no module was found for computing representative cycles.
- **`Ripser`:** A popular method written in C++ [46]. There is experimental support for computing representative cycles (in a separate branch).
- **`Ripser.py`:** builds on top of `Ripser` with computations of representative cocyles. As it is built on `Ripser` it might be possible to also get representative cycles, however it was not trivial.

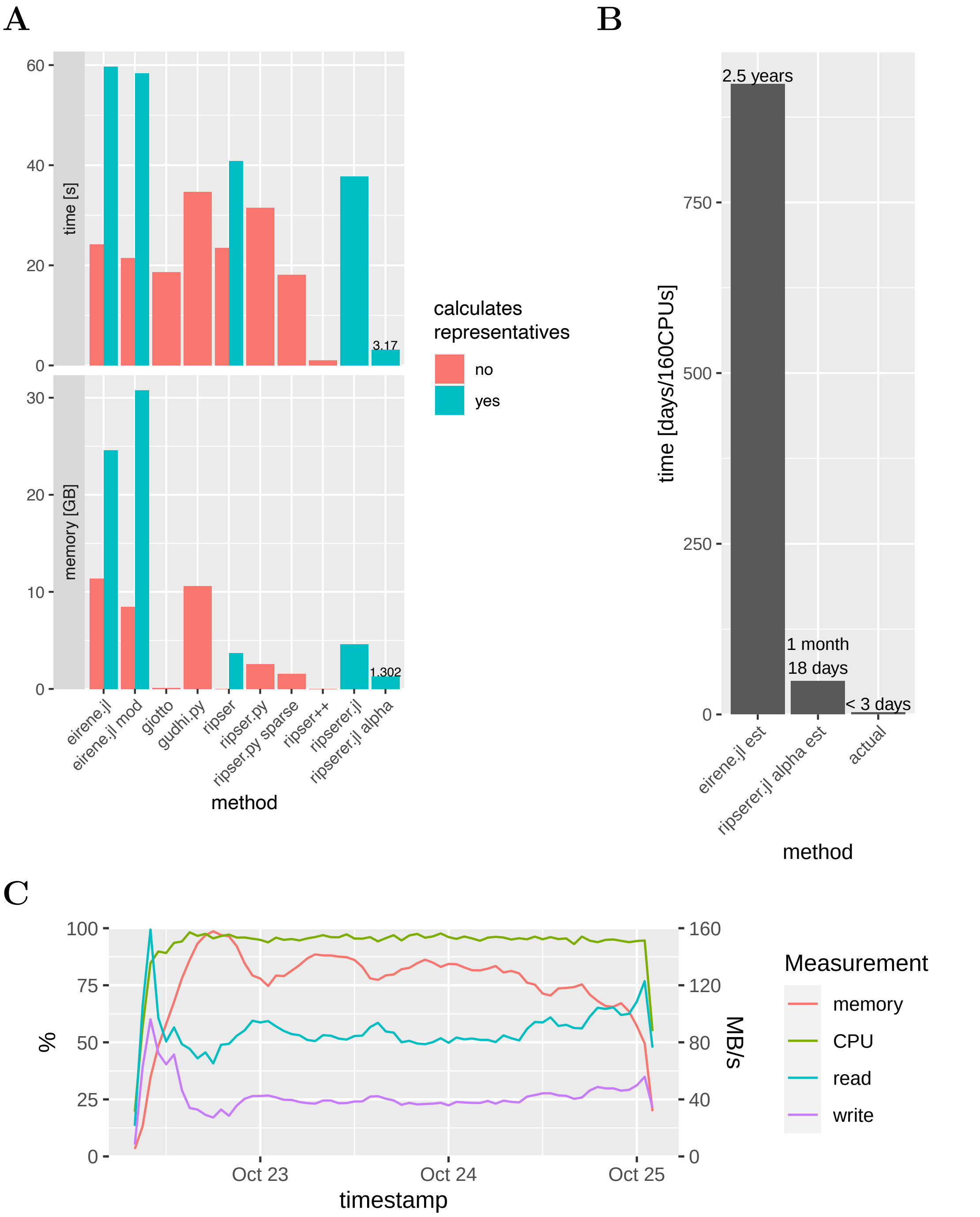


**Supplementary Figure 4. Performance benchmarks.** Performance comparison for a number of tested persistent homology tools (A). Bar fill indicates if representative cycles were included in the output. All benchmarks were on a single large structure with 600 residues. Actual computational on all 214M structures compared to estimates based on the benchmark of the initial method and the chosen method (B). Percentage use of the 1 TB memory and 160 CPUs, as well as disk read/write in MB/s during the 3 day computation (C).

`Ripser.py sparse:` an approximate sparse filtration with sparse distance matrix tested to reduce computational time.

**`Ripser++`:** The only GPU method tested [54]. Clearly this is a big advantage, however it was not possible to compute representative cycles.

**`Ripserer.jl`:** a Julia implementation of `Ripser` [25].

**`Ripserer.jl alpha`:** by default, `Ripserer.jl` (and all other listed tools) uses Vietoris-Rips filtration (Fig 1). Alpha filtration was tested here, which can be much more efficient on low-dimensional point clouds.

Computational time for `Eirene.jl` and `Ripserer.jl` with Alpha filtration were estimated simply by multiplying computational time observed in Fig 4A by 214M and dividing by 160 (Fig 4B). The runs are assumed to be completely parallel since multiple identical calls to `Ripserer.jl` will be performed each given a single core. It is a rough estimate since the average number of residues is around 333, however computational time does not scale linearly with residue count; larger point clouds take up a disproportionate amount of the total time. The estimated time was more than 16 times longer than the actual. This is partly explained by the large point-cloud used for benchmarking, essentially making it a worst case estimate, and partly explained by a few other optimisations:

**TAR iteration:** Instead of extracting and reading files in the sharded proteomes it was found to be much more efficient to stream the content of the TAR archives directly using `TarIterators.jl` (with a minor tweak).

**CIF parsing:** Instead of reading the CIF files with a standard CIF reader they were instead streamed line-by-line only reading a required subset of the file contents.

**Centrality on sparse $H$:** The hypergraph centrality code was rewritten and tailoured to this project's specific use-case, particularly with a sparse representation of the hypergraph $H$, paired with an efficient implementation of the sparse encoding itself.

The output of the topological analyses was written to compressed JSONs matching the structure of the shared proteomes, and later repackaged into HDF5 files to organise by UniProt accessions [32], to allow for partial read/write and in order to add additional protein meta-data.

### 2. The topological tree of life

The taxonomy tree is visualised in Fig 1 of the main text with a circle packing plot, is generated by constructing circles for each species with area proportional to its number of AlphaFold2 structures (including any entries annotated with its subspecies and other lower ranks). Circles associated with child nodes of genuses are then circle packed, one genus at the time. This process is repeated for each rank going up, which means that the area of higher ranks is only approximately representative of their number of structures.

The lightness of circles indicates the *topological richness* of the proteins belonging to a taxonomy ID. The richness is defined as the persistence of the 1-dimensional topological features, restricted to those having persistence $\geq 10$, divided by the number of residues in the protein and averaged across proteins.

Each edge in the taxonomy tree is represented visually as outlines around the circles. The outlines are sized according to the taxonomy rank, with slightly thinner outlines for lower ranks. The outlines are colored in a black-to-yellow palette, indicating the variance of the number of 1-dimensioanl topological feature in each protein, normalized by the number of proteins in the circle. Differences in the outlines are made clearer by a log-transform, specifically $\log_{10}$ of one minus the correlation. To check whether the variance was influeced by the number of residues in each protein, we further normalized by this quantity. The output of this latter computation has a 0.925 correlation coefficient with the non-normalized one, showing thus high consistency.

Circles are packed within each container circle with the R library `packcircles` [52] and visualized with `ggplot2` [53]. Data is from the tables `TreeNode` and `TreeEdge` from the Postgres database, as well as the table `AF` for the zoomed example for Hemoglobin.

### 3. Mediaflux

We share the output of our topological analysis on Mediaflux. Here, the data is organized into three folders: compressed JSON files, HDF5 files, and a Postgres database. The entire datasets can be downloaded with the following links: JSON (∼10 TB), HDF5 (∼9 TB), and Postgres database (∼210 GB). The links will not immediately start downloads but rather prompt for installing a helper utility "Mediaflux Data Mover" which will then aid in the download process.

See Fig. 5 for an overview of the data structure. Some data containers are left blank for simplicity.

#### 3.1. **JSON.**

Protein structures predicted by AlphaFold2 and topological data is stored in GZip compressed JSONs. The organization is similar to the proteome sharing provided by AlphaFold2. Additionally, sharded proteomes are placed in folders according to the first three numbers of the taxonomy id. The JSONs contain integers and floats (floating-point values). Numbers are either provided as scalar, in a list or lists of lists. Newline in the figure indicates the highest grouping level for JSON values.

**n:** number of residues (scalar).

**x, y, z:** α-carbon coordinates in Å(list of floats).

**cent1, cent2:** TIFs for dimension 1 and 2 (list of floats).

**bars1, bars2:** Birth and death filtration times for each topological feature in dimension 1 and 2 (list of floats).

**reps1, reps2:** Representative cycles for dimension 1 and 2. Stored as a list of lists of integers. Each representative cycle is a set of either 1- or 2-simplices, provided as node indices (1-indexed).

For each proteome, we also include the topological clusters computed as graph-communities, as explained in [26] and using Python's Louvain module [45]. The result is written to a compressed JSON with one entry per accession, containing community indexes for each residue.

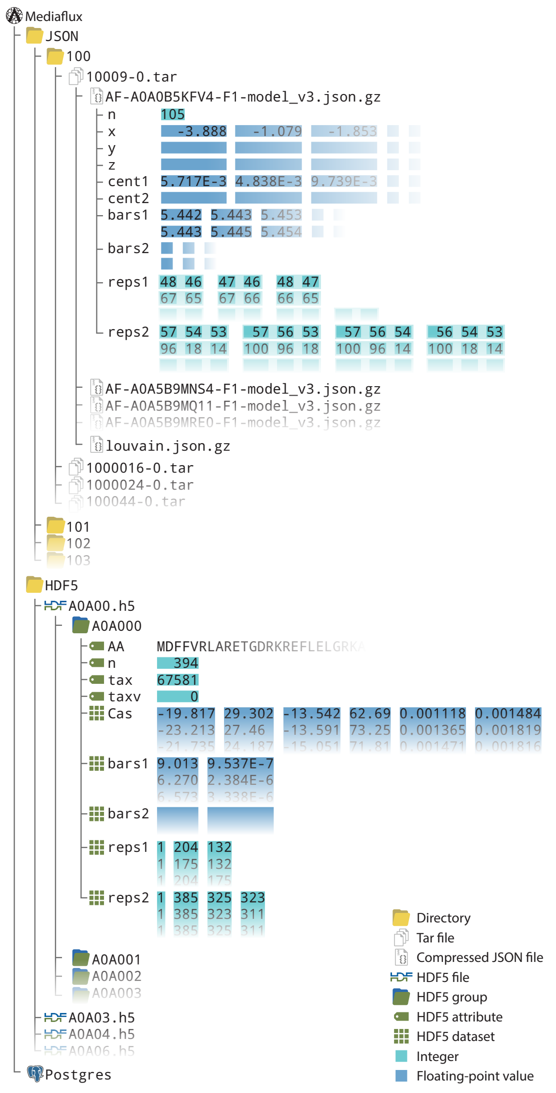


**Supplementary Figure 5. Data organization on Mediaflux.** One example protein is shown for JSON format as well as HDF5.

#### 3.2. **HDF5.**

The data is also provided in Hierarchical Data Format version 5 (HDF5) organized by UniProt accession. Proteins are placed together in HDF5 files based on the first five characters of their accessions. Each protein is found as a HDF5 group, which contain HDF5 attributes and HDF5 datasets. Here, each dataset is always a table of unnamed columns, stored as a numerical matrix.

`AA`: One-letter amino acid sequence encoded as an ASCII string.
`n`: Number of residues.
`tax`, `taxv`: Taxonomy ID and sharding index used by AlphaFold2.
`Cas`: Values for each node, i.e. α-carbons. The columns are the x, y, z coordinates in Å, pLDDT score (AlphaFold2 confidence score), and TIFs in dimension 1 and 2.
`bars1`, `bars2`: Birth and persistence (death − birth) filtration times for each topological feature in dimension 1 and 2.
`reps1`, `reps2`: Representative cycles for dimension 1 and 2. The first column is an index for the feature, starting at 1. The remaining columns are node indexes for members of a simplex (one simplex per row).

**Remark** (Decompression step needed to access files)**.** The HDF5 files are uncompressed except for the datasets `reps1` and `reps2`, which require a ZStd plugin (Zstandard) for access. For example, in python `import h5py, zstandard` and in Julia `using HDF5, H5Zzstd` will suffice to read the compressed datasets.

3.3. **Postgres.** Protein metadata is collected in a Postgres database 6.

***AF*** 214,175,790 × 30

| acc | taxon | n | maxrep1 | maxrep2 | maxpers1 | maxpers2 | meanplddt | nrep1 | nrep2 | nrep1_t1 | nrep1_t2 |
|---|---|---|---|---|---|---|---|---|---|---|---|
| A0A4Y2VRE5 | 182803 | 143 | 74 | 36 | 4.869422 | 2.108323 | 80.50973 | 310 | 189 | 27 | 4 |
| A0A4Y2VRE6 | 182803 | 200 | 24 | 46 | 3.9872656 | 1.7559824 | 77.5748 | 390 | 283 | 19 | 8 |
| A0A4Y2VRE9 | 182803 | 104 | 5 | 6 | 0.67140913 | 0.15324402 | 88.001144 | 198 | 167 | 0 | 0 |

***Tax*** 888,163,507 × 2

| acc | tax |
|---|---|
| A0A009IHW8 | 1310613 |
| A0A011QK89 | 522306 |
| A0A017SE81 | 1388766 |

***JSON*** 214,175,790 × 2

| acc | path |
|---|---|
| A0A3S0EAG4 | PH/100/100-0/AF-A0A3S0EAG4-F1-model_v3.json.gz |
| A0A3S0EAU0 | PH/100/100-0/AF-A0A3S0EAU0-F1-model_v3.json.gz |
| A0A3S0EB17 | PH/100/100-0/AF-A0A3S0EB17-F1-model_v3.json.gz |

***TaxTree*** 4,367,074 × 5

| tax | parent | rankp | rank | domain |
|---|---|---|---|---|
| 3040113 | 3040113 | species | species | V |
| 3040114 | 3040114 | species | species | V |
| 3040115 | 3040115 | species | species | V |

***TaxParent*** 47,595,713 × 4

| tax | parent | rankp | rank |
|---|---|---|---|
| 1211725 | 2626993 | no rank | species |
| 1211726 | 264000 | genus | species |
| 1211727 | 2646371 | no rank | species |

***TreeNode*** 879,207 × 40

| tax | domain | proteins | avg_n | avg_maxrep1 | avg_maxrep2 | avg_maxpers1 | avg_maxpers2 | avg_nrep1 | avg_nrep2 | var_n |
|---|---|---|---|---|---|---|---|---|---|---|
| 1955042 | B | 2758 | 294.74 | 109.33828861 | 38.09499637 | 6.612734582391 | 1.8536213896 | 711.630529 | 463.369108 | 39083. |
| 1956187 | E | 14966 | 365.27 | 118.60216784 | 38.7913651163 | 10.6211077741 | 1.84818517341 | 822.681202 | 531.995778 | 61255. |
| 1961800 | E | 11 | 322.15 | 98.7333333333 | 52.755555555 | 6.632029215494 | 2.00836224026 | 699.222222 | 383.955555 | 837.62 |

***TreeEdge*** 871,760 × 17

| child | parent | proteins_f | avg_n_f | avg_maxpers1_f | avg_maxpers2_f | avg_nrep1_f | avg_nrep2_f | var_n_f |
|---|---|---|---|---|---|---|---|---|
| 2039065 | 30080 | 0.000431220 | 0.4838341 | 0.20544891606081 | 0.07054937392533 | 0.29337834246 | 0.35278577095 | 0.0000000 |
| 2039066 | 30080 | 0.001293661 | 1.0048862 | 1.05280799547665 | 0.96839498613992 | 0.96395741096 | 0.93714374028 | 2.0785838 |
| 2039094 | 30080 | 0.000431220 | 1.4639083 | 1.3356359567923 | 2.093408533313980 | 1.64743223078 | 1.48712771141 | 0.0000000 |

**Supplementary Figure 6. Example entries from database tables.** The first three rows to exemplify the data contained in the protein metadata. Table sizes indicated with rows × columns notation.

`AF`: The main table, which contains summary statistics computed on the topological analysis results for each protein.
`JSON`: Path to JSON file for a given UniProt accession.
`Tax`: Taxonomy ID associated with a vast amount of identifiers (NCBI taxonomy FTP server).
`TaxTree`: Taxonomy ids at species level or lower, with species parent indicated. Species as child nodes are also included with themselves as parent. This table (in combination with `Tax`) is used for connecting any relevant accession to a species.
`TaxParent`: All direct and indirect child nodes for a subset of taxonomy ranks (Domain, Kingdom, Phylum, Class, Order, Family, Genus, and Species).

**TreeNode:** Taxonomy tree nodes with summary statistics for the same subset of taxonomy ranks as in TaxParent.

**TreeEdge:** Taxonomy tree edges and summary statistics between the nodes from TreeNode.

Fig 1 in the main text is build from the tables TaxNode and TaxEdge, after further data processing (see data/alphafold/vis/ in the code repository).

**Supplementary Table 1. Database schema.** The description of each column of each table in the database.

| Table | Column | Type | Description |
|---|---|---|---|
| AF | acc | string | UniProt accession ID |
| AF | taxon | integer | Taxonomy ID used by AlphaFold2 |
| AF | n | integer | Number of residues |
| AF | maxrep1 | integer | Maximum number of residues in a feature (dim=1) |
| AF | maxrep2 | integer | Maximum number of residues in a feature (dim=2) |
| AF | maxpers1 | float | Maximum persistence of a feature (dim=1) |
| AF | maxpers2 | float | Maximum persistence of a feature (dim=2) |
| AF | meanplddt | float | pLDDT averaged across α-carbons |
| AF | nrep1 | integer | Number of representatives (dim=1) |
| AF | nrep2 | integer | Number of representatives (dim=2) |
| AF | nrep1_t$T$ | integer | nrep1 with persistence $> T$ ($T \in [1, 10]$) |
| AF | nrep2_t$T$ | integer | nrep2 with persistence $> T$ ($T \in [1, 10]/10$) |
| JSON | acc | string | UniProt accession ID |
| JSON | path | string | Path in database to JSON file |
| Tax | acc | string | UniProt accession ID |
| Tax | tax | integer | Taxonomy ID |
| TaxTree | tax | integer | Taxonomy ID for species and lower ranks |
| TaxTree | parent | integer | Taxonomy ID for species |
| TaxTree | rankp | string | "species" |
| TaxTree | rank | string | Rank of tax ("species" or lower) |
| TaxTree | domain | string | Domain of tax ("A", "B", "E", or "V") |
| TaxParent | tax | integer | Taxonomy ID |
| TaxParent | parent | integer | Taxonomy ID for direct or indirect parent of tax |
| TaxParent | rankp | string | Rank of parent |
| TaxParent | rank | string | Rank of tax |
| TreeNode | tax | integer | Taxonomy ID |
| TreeNode | domain | string | Domain ("A", "B", "E", or "V") |
| TreeNode | proteins | integer | Number of proteins |
| TreeNode | avg_$V$ | float | AF values $V$ averaged across child nodes |
| TreeNode | avg_$V$_pp | float | AF values $V$ averaged across proteins |
| TreeNode | var_$V$ | float | Variance of AF values $V$ averaged across child nodes |
| TreeNode | var_$V$_pp | float | Variance of AF values $V$ |
| TreeEdge | child | integer | Child node taxonomy ID |
| TreeEdge | parent | integer | Parent node taxonomy ID |
| TreeEdge | $V$_f | float | $V_{\text{child}}/V_{\text{parent}}$ |

### 4. Datasets and Supplementary Results

The datasets discussed in the results are summarized in Table 2. In each of these datasets, we removed structures with low-confidence AlphaFold2 predictions. AlphaFold2 produces a per-residue confidence score (pLDDT) [2], which assigns a value between 0 and 100 to each residue in a structure; values below 70 are considered low. Here, to select proteins with an overall good prediction, we average the pLDDTs over all the residues in a structure, and discard those scoring an average below 70. The remaining ones are considered *high confidence* predictions, and are kept in the dataset.

**Supplementary Table 2. Dataset overview.** A brief summary of the datasets discussed in the biological analysis. We analyzed all the AlphaFold2 structures and in the biological comparisons focused on the high confidence structures. More detailed discussions of each data set are given in the text below.

| Dataset | AlphaFold2 | High confidence |
|---|---|---|
| RCSB | 2712 | 2637 |
| M-CSA | 866 | 862 |
| CATH | 73749 | 62861 |
| Thermo-/mesophiles [51] | 1815 | 1656 |
| Mutations(disease/neutral) | 471/855 | 419/681 |

4.1. **Comparison with experimental structures (RCSB).** To compare between the topological analysis performed on AlphaFold2 predictions and on experimental structures, we considered all the 2,712 UniProt entries with full structure available on PDB. These UniProt accessions correspond in total to 28,309 different experimentally solved protein chains. Out of the 2,712 AlphaFold2 predictions, only 2,637 have a high-confidence score. For each of these structure, we considered the 1 and 2-dimensional TIFs, and computed the correlation coefficient between the resulting vector for each predicted structure and its experimental counterparts.

Complete lists of the structures considered in each dataset, and the correlation coefficients, are available for download, see Data Availability. This folder contains:

- a file `uniprot2PDB_fullstructures.json`, containing mapping between UniProt accessions and PDB entries
- a file `centrality_correlation.csv`, containing, for each UniProt id, the correlation coefficient between its 1 and 2-dimensional topological influence vectors and the experimental counterparts

Correlation coefficients were computed using `numpy`'s `corrcoef` function.

4.2. **M-CSA dataset.** To analyze the relation between 2-dimensional topological clusters and binding sites, we looked at the Mechanism and Catalytic Site Atlas (M-CSA) [36], a database of enzyme reaction mechanisms, which provides catalytic residues of hundreds of enzymes. We downloaded all the 1033 PDB entries of experimental structures with annotated sites, and performed the topological analysis on the corresponding structures, see Figure 7 for the result of our analysis.

To reproduce the result on AlphaFold2 predicted structures, we then mapped each PDB entry to the corresponding UniProt accession, when found. This left us with a total of 866 different proteins, 862 of which are predicted by AlphaFold2 with a high

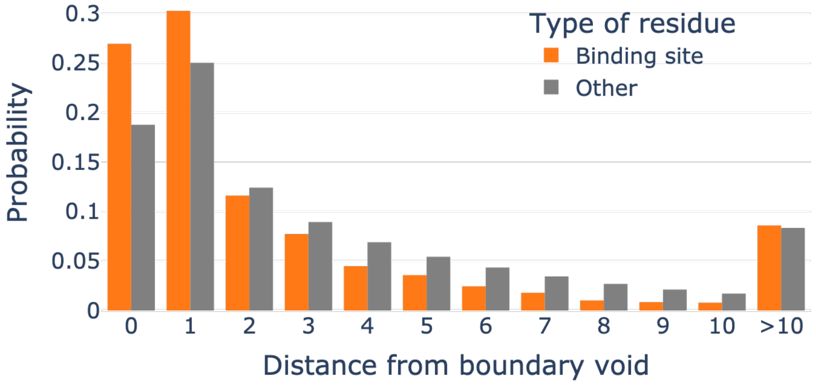


**Supplementary Figure 7.** The bar plot shows the distribution of distances (in number of residues) of residues between binding sites from boundaries of 2-dimensional topological clusters in experimentally solved structures. These boundary points are enriched for binding sites from the Mechanism and Catalytic Site Atlas (M-CSA) dataset.

confidence score.

A complete list of the structures considered in each dataset, and code to reproduce the results, are available for download, see Data Availability. This folder contains:

- a file `CSA_site.tsv`, containing PDB entries and residue numbers of binding sites
- a file `CSA_AF.csv`, containing mapping between PDB and UniProt accessions, as well as the confidence score of the AlphaFold2 predictions
- files `communities.json` and `communities-experimental.json` containing the partition of each structure into 2-dimensional topological clusters. The organization of these JSON files is as described in Section 3.1
- notebooks `Results.ipynb` and `Results_experimental.ipynb` to compute boundary points between 2-dimensional topological clusters and to reproduce the results

4.3. **CATH.** To investigate the relation between topological features and protein domains, we looked at all the 73,749 AlphaFold2 predictions containing at least two distinct identified CATH domains [31]. These structures, and the corresponding domain mapping, were recently identified in [24]. We then excluded low-confidence predictions (leaving 62,861 proteins) and reduce the dataset to a list of 38,171 non-redundant structures. This last step was achieved using the software `CD-HIT` [49] and a threshold of 70% sequence similarity.

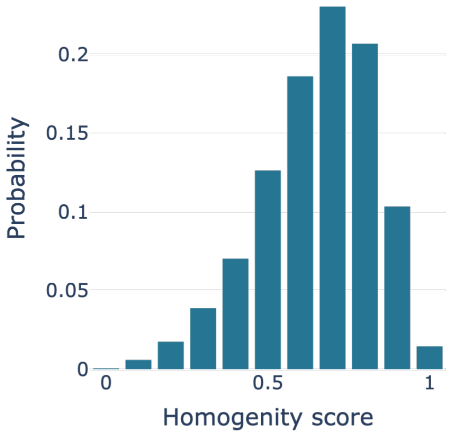


**Supplementary Figure 8.** The bar plot shows the distribution of homogeneity scores for all the 62,861 high-confidence AlphaFold2 predictions with identified CATH domains.

To quantify the agreement between the partition induced by CATH domains and by 1-dimensional topological cluster we computed the homogeneity score using the `homogeneity_score` function in Python's `sklearn` package. The homogeneity score is a value between 0 and 1; a clustering satisfies homogeneity (and thus has homogeneity 1) if all of its clusters (in our case 1-dimensional topological clusters) contain only data points which are members of a single class (in our case, a single CATH domain). For completeness, Figure 8 shows the results for the 62,861 high-confidence AlphaFold2 predictions, including redundant ones.

A complete list of the considered structures, the corresponding homogeneity scores, their partition into 1-dimensional topological clusters and CATH domains are available for download, see Data Availability. This folder contains:

- a `hom_scores_red.csv`, with UniProt entries, homogeneity score, confidence score of the prediction, and whether they are non-redundant or not
- a file `domain_vectors.json` with the partition into CATH domains
- a file `communities_all.json` with 1-dimensional topological clusters

4.4. **Thermophiles and mesophiles.** To investigate structural differences between enzymes in thermophilic and mesophilic organisms, we selected 10 different Enzyme Commission (EC) numbers based on their biotech relevance, a total of 30 thermophilic and 8 mesophilic organisms, and we listed all UniProt entries with these characteristics. In total, we considered 1,815 different protein structures, that became 1,656 after excluding low-confidence predictions. On this latter dataset, we were interested in analyzing the distribution of volumes of significant 2-dimensional features (*i.e.* with high persistence).

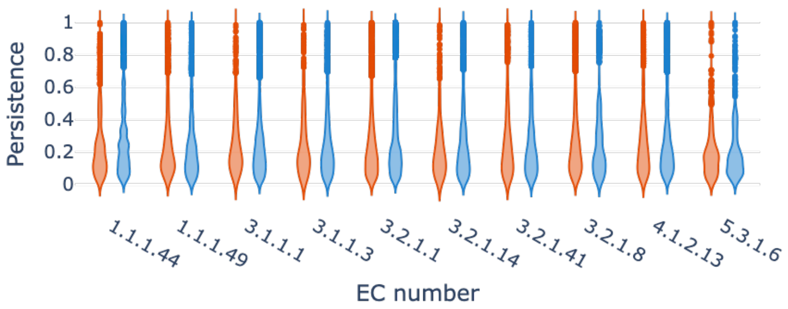


**Supplementary Figure 9.** The distribution of persistence values of 2-dimensional topological features across the different EC numbers, for low-persistence ($< 1$) features. Thermophiles are in red and mesophiles in blue.

The distribution of features with persistence $< 1$ turned out to be almost identical across EC numbers and thermal characteristics, see Figure 9. For this reason, we restricted our attention to topological features with persistence $\geq 1$, that show more variation, see Figure 10.

The volume of each feature was computed using `scipy ConvexHull` function, as the volume of the convex-hull of residues in the generator. Our results show that mesophile organisms have on average larger voids in their enzymes, and that this patter is robust. In Figure 11, error bands are given by sampling 1000 different voids in thermophiles and mesophiles, respectively, and then looking at the standard deviation.

A natural question is whether this pattern is maintained for single EC numbers. Volume, number, and persistence of voids are all strongly influenced by the size and length of the protein. Since the distribution of lengths in individual EC numbers is different for thermophiles and mesophiles, to analyze EC number, we first selected thermophilic and mesophilic proteins in the same range of length. For a given EC number, this is achieved by randomly selecting a mesophilic enzyme for each thermophilic one, with a difference in length of at most 5 residues. The result of this analysis are shown in Figure 12.

A complete list of the structures considered, and code to reproduce the results, are available for download, see Data Availability. This folder contains:

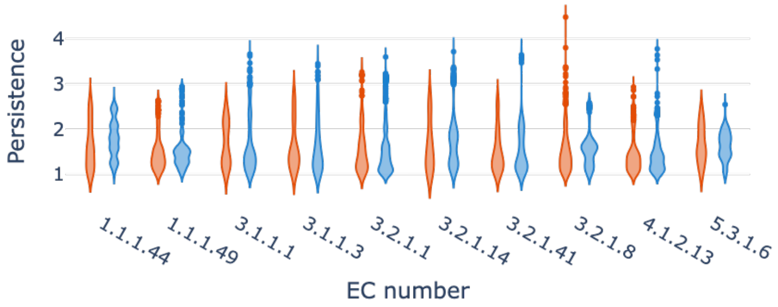


**Supplementary Figure 10.** The distribution of persistence values of 2-dimensional topological features across the different EC numbers, for high-persistence ($\geq 1$) features. Thermophiles are in red and mesophiles in blue.

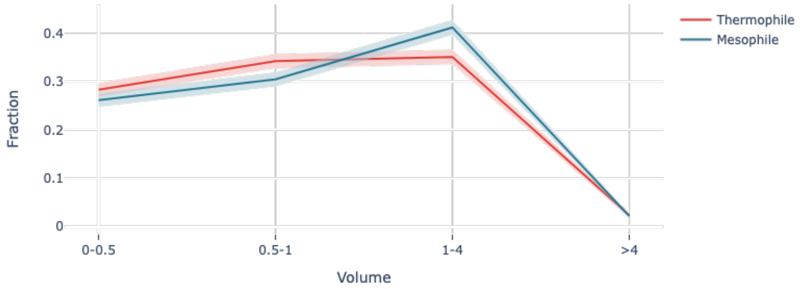


**Supplementary Figure 11.** The distribution of volumes of 2-dimensional topological features, for high-persistence ($\geq 1$) features. Error bands are given by standard deviation. Thermophiles are in red and mesophiles in blue.

- a file `thermozymes-acc-unjag-summ.tsv`, containing accessions and taxonomy information of the structures considered
- a file `summary.csv`, containing confidence scores of the structure considered
- a file `thermo_all.csv` containing the volumes and persistence values of the 2-dimensional topological features
- a file `SEQ.csv` containing TIFs for different amino acids
- a file `Samples.csv`, containing the distribution of volumes for the sampled dataset
- a notebook `Results.ipynb` containing code to reproduce the sampling used for the result in Figure 12

4.5. **Mutations.** To check if our analysis is effective in the detection of protein regions that are enriched for damaging mutations, we looked at the datasets of disease-causing and neutral variants studied in the paper [43], where the authors consider a few hundreds experimental structures and their disease-associated missense variants, and link damaging mutations to structurally damaging changes in their mutant structures.

As usual, we restrict, our analysis to structures with high-confidence predictions. Results in the manuscript show the distribution of 2-dimensional TIFs for residues

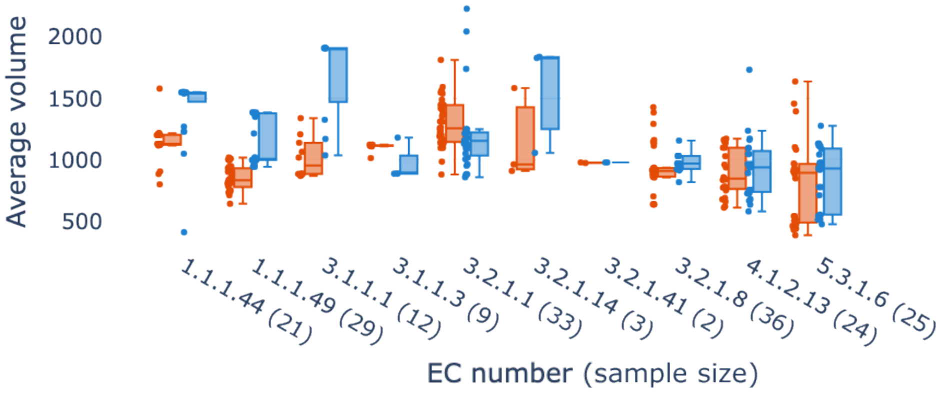


**Supplementary Figure 12.** The distribution of average volumes of 2-dimensional topological features across the different EC numbers, for high-persistence ($\geq 1$) features. The pattern is maintained across all but two groups. Thermophiles are in red and mesophiles in blue. Sample sizes in parentheses are numbers of proteins for thermophiles plus mesophiles.

**A** Main dataset **B** Control group

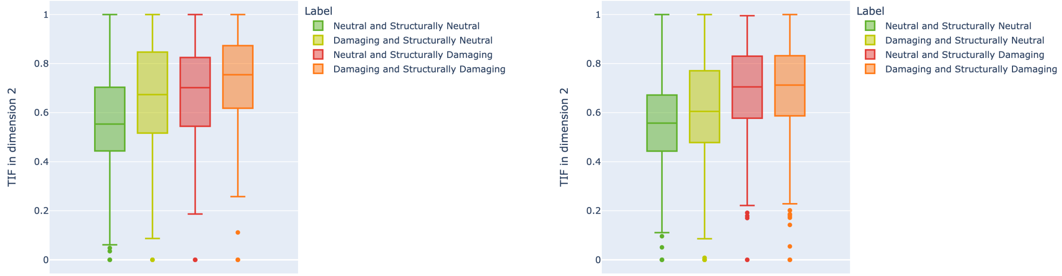


**Supplementary Figure 13.** The distribution of 2-dimensional TIFs in residues accommodating missense variation considered in [43].

accommodating neutral mutations that do not cause structural damage, and disease-associated mutations that modify the structure. Figure 13A shows the distributions for the full set of labels, and Figure 13B shows the same result for the control dataset used in [43]. As shown in Figure 14, the pattern is maintained for 1-dimensional TIFs, although the differences are weaker.

The data for the ACE2 and HBB examples shown in the manuscript is taken from the Missense3D database [44], which catalogs amino-acid substitutions that are predicted to be structurally damaging [43, 44]. A third example we analyzed is CFTR, the results are shown in Figure 15.

A complete list of the structures considered is available for download, see Data Availability. This folder contains:

**A** Main dataset **B** Control group

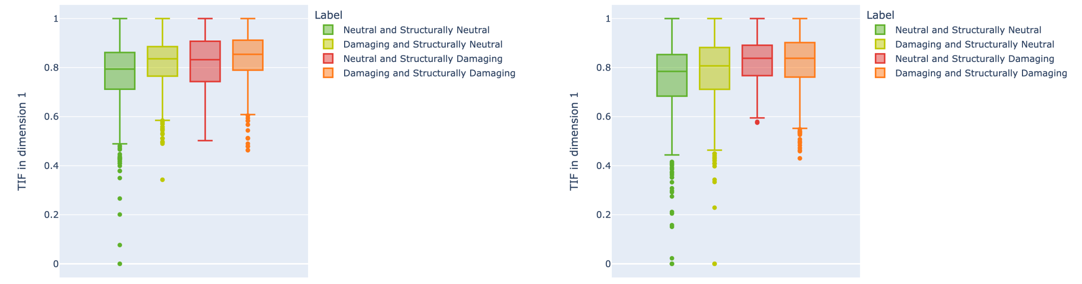


**Supplementary Figure 14.** The distribution of 2-dimensional TIFs in residues accommodating missense variation considered in [43].

- files `mutations.csv`, `mutations_control.csv`, containing UniProt accessions of the proteins considered and the list of mutations with labels and TIFs;
- files `ace2_cent.csv`, `cftr_cent.csv`,`hbb_cent.csv`, containing the list of mutations with labels and TIFs for the examples shown;
- a file `thermo_all.csv` containing the volumes and persistence values of the 2-dimensional topological features;
- a notebook `Results.ipynb` containing code to visualize the results.

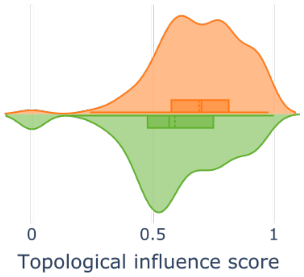


**Supplementary Figure 15.** The distribution of 2-dimensional TIFs for residues in human CFTR that are predicted to accommodate structural damaging mutations.

**Data Availability.** The datasets analyzed for the current study are available for direct download from MediaFlux via: https://bit.ly/protTDA (1.31 GB). All project code is available in the protTDA repository: https://github.com/degnbol/protTDA. Topology outputs can be bulk downloaded from MediaFlux: JSON files via https://bit.ly/protTDAjson (∼10 TB), and HDF5 files via https://bit.ly/protTDAhdf5 (∼9 TB). The Postgres database containing protein and taxonomy data is available via https://bit.ly/protTDAdb (∼210 GB).

[53] Hadley Wickham. *ggplot2: Elegant Graphics for Data Analysis*. Springer-Verlag New York, 2016.

[54] Simon Zhang, Mengbai Xiao, and Hao Wang. Gpu-accelerated computation of vietoris-rips persistence barcodes. In *36th International Symposium on Computational Geometry (SoCG 2020)*. Schloss Dagstuhl-Leibniz-Zentrum für Informatik, 2020.
